# Computational quantification of global effects induced by mutations and drugs in signaling networks of colorectal cancer cells

**DOI:** 10.1101/2020.12.30.424842

**Authors:** Sara Sommariva, Giacomo Caviglia, Silvia Ravera, Francesco Frassoni, Federico Benvenuto, Lorenzo Tortolina, Nicoletta Castagnino, Silvio Parodi, Michele Piana

## Abstract

Colorectal cancer (CRC) is one of the most deadly and commonly diagnosed tumors worldwide. Several genes are involved in its development and progression. The most frequent mutations concern APC, KRAS, SMAD4, and TP53 genes, suggesting that CRC relies on the alteration of different pathways. However, with classic molecular approaches, it is not easy to simultaneously analyze the interconnections between these pathways. For this reason, we propose a computational model based on a huge chemical reaction network to simulate the effects induced on the global signaling associated with CRC by single or multiple concurrent mutations or by drug treatment. This approach displays several advantages. The model can quantify the alteration in the concentration of the proteins connected with the examined mutation. Moreover, working on the global signaling of CRC, it is possible to disclose unexpected interactions between the involved pathways, representing new therapeutic targets.

**Highlights:**

1. Colorectal cancer relates to defects in many different pathways within cell signaling
2. Cell signaling is modeled as a chemical ration network with 10 interacting pathways
3. Global effects induced by single or multiple concurrent mutations are quantified
4. A possible extension of the model to account for a targeted drug is discussed

## Introduction

Colorectal cancer (CRC) is the second most common cancer in women (823,303 estimated incident cases worldwide, according to the GLOBALCAN 2018 data, https://gco.iarc.fr/) and the third most common in men (1,276,106 estimated incident cases worldwide) (Armaghany et al., 2012; Tariq and Ghias, 2016), representing one of the most significant causes of cancer death (Rawla et al., 2018). Both genomic and epigenetic alterations are common in CRC and are the driving forces of tumorigenesis.

CRC arises from one or more of three mechanisms: the chromosomal instability pathway, the microsatellite instability pathway, and the CpG island methylator phenotype. Focusing the attention on the chromosomal instability pathway, since 1990, it is well known that the mutations of genes involved in cell growth and differentiation pathways play a pivotal role in the development and progression of CRC (Fearon and Vogelstein, 1990). Although numerous mutations have been identified associated with the CRC development and progression, the most frequent driver and gate-keeper mutations commonly found concern TP53, APC, KRAS, PTEN, SMAD4, PIK3CA, BRAF, AKT (Anderson et al., 2019; Armaghany et al., 2012; Castagnino et al., 2016; Smith et al., 2002; Tariq and Ghias, 2016; Tortolina et al., 2015). In addition, several indications suggest that the aggressiveness and malignancy of CRC depend on the mutation order. In particular, it has recently been observed that the most frequent mutation order in CRC is the following: APC, KRAS, SMAD4, and TP53 (Levine, 2019).

Since these genes belong to distinct signaling pathways, it may be concluded that many CRCs should be related to alteration in the normal communications between interconnected pathways. According to these findings CRC may be seen as a multifaceted disease, while the study of single pathways might be not sufficient to individuate the principal effectors responsible for its proliferation and malignancy (Logue and Morrison, 2012; Lun and Bodenmiller, 2020; Sever and Brugge, 2015).

Intracellular changes induced by mutations have been investigated at different levels (Creixell et al., 2015). A very basic and natural one is the interaction between proteins to form complexes. This approach typically involves a huge number of interactions but is not sufficient to fully understand the changes induced by a mutation in the behavior of a complex system of chemical reactions. On a slightly higher scale, an alternative, functional level is based on grouping proteins in pathways, essentially designed to achieve a defined cellular process (Lin et al., 2007), which is often visualized through a linear diagram. Pathways may be of help in explaining the role of a protein or its possibly mutated form, but a protein can participate in two or several pathways, thus affecting more cellular functions, which in turn may be influenced by other mutations. Accordingly, interconnections, superpositions, and nonlinearities may put severe limitations on the insight provided by the use of pathways (Khatri et al., 2012).

Here we follow a synthetic view, whereby we consider a set of proteins whose mutated forms are frequently observed in CRC, and we regard them as particular nodes of a large intracellular signaling network. Signal transmission is accomplished by a cascade of reactions organized into a chemical reaction network (CRN), henceforth denoted as CRC-CRN.

Specifically, the starting point of the present analysis is the CRC-CRN modeling the intracellular processing of the information sensed from the environment through the TGFβ, WNT, and EGF families of receptor ligands, at the G1/S transition point of healthy colorectal cells (Castagnino et al., 2016; Tortolina et al., 2015). This results in a complex CRN, that comprises hundreds of proteins and tens of reaction pathways with associated ramifications.

By applying mass action kinetics and using standard procedures, the CRC-CRN is mapped into a system of ordinary differential equations (ODEs), thus allowing for simulations of the kinetics of the signaling process (Anderson et al., 2019; Chellaboina et al., 2009; Roy and Finley, 2017). The resulting mathematical model is capable of describing the behavior of healthy physiological networks, as well as the consequences of individual pathological conditions associated with specific mutations. As for mutations, we adopt the formal description of Loss of Function (LoF) and Gain of Function (GoF) that has been extensively analyzed in a previous paper (Sommariva et al., 2020).

In the present work, starting from the system of ODEs, we want to develop an efficient computational tool that can be used for an in silico analysis of mutations of the CRC-CRN and of the effects induced by targeted drugs.

## Results

### The CRC-CRN as a simulation tool for biological analysis of cancer cells

This study was performed focusing our attention to the G1-S transition phase of CRC cells. We have considered a relevant sub-region of the cell’s signaling network downstream the constant external growth factors TGFβ, Wnt, and EGF, which are all involved in CRC (Koveitypour et al., 2019). This has provided the appropriate mathematical framework to simulate the global and quantitative effects of the LoF and GoF mutations most frequently accumulated in CRC cells.

The present approach is inherently global, in that it is capable of considering the combined effects of chemical reactions involving several proteins belonging to the physiological signaling network, as well as of describing the overall changes induced by mutations in a cancer cell. The network provides a comprehensive view of the cascade of reactions, examining them not as components of a single pathway, but in a global and quantitative way. In particular, the mathematical model associated with the CRN provides the equilibrium values of the concentrations of the proteins involved in the network and enables to compare the physiological values with those induced by single or multiple mutations. To this end, the concentrations of the 419 proteins within the CRC-CRN are regarded as the state variables of a dynamical system of 419 ODEs, which describe 850 biochemical reactions following the mass action law and thus depending on as many rate constants. The stable equilibrium states of the network are identified with the stationary states of the system, achieved at large time values; they are computed by numerical integration of the ODEs of the model, after setting the initial values of the protein concentrations.

In principle, simulations through dynamical systems imply dependence of the results on the initial conditions, which in our case consist of a set of 419 parameters describing the values of the protein initial concentrations. As shown in the Model and Methods Section, the number of the required input parameters actually reduces to 81. Indeed, every equilibrium state is uniquely defined by setting the values of the constant aggregation concentrations within the moiety conservation laws of the network or, equivalently, by fixing a stoichiometric compatibility class.

Mutations resulting in the LoF or GoF of a selected protein have been incorporated in the CRC-CRN as described in the Model and Methods section and synthetically depicted in Figure 1. By computing the concentrations at the equilibrium of the resulting modified network, we are able to rigorously quantify the impact of the considered mutations on the whole set of proteins involved in the CRC-CRN.

**Figure 1:**
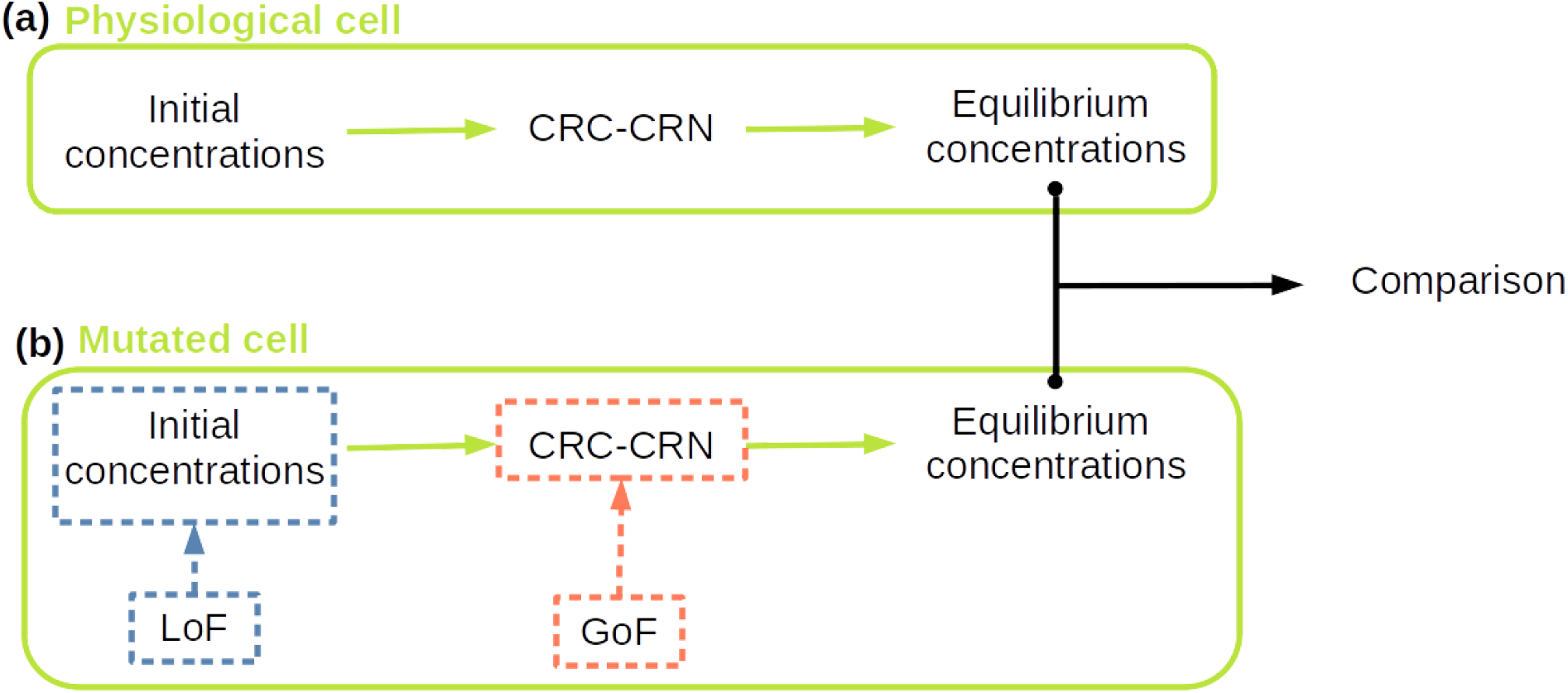
General workflow of the CRC-CRN. (a) The values of the protein concentrations at the equilibrium in the physiological cell are computed by integrating the system of ODEs defining our CRC-CRN. (b) Any mutation resulting in the loss of function (LoF) of a protein is embedded in our model by modifying the values of the conserved moiety in input. Conversely, mutations causing gain of function (GoF) are implemented by modifying the system of ODEs. The asymptotically stable state of the modified system defines the protein concentrations at the equilibrium in the mutated cell.

### Simulation of global effects induced by a single-gene mutation

To evaluate whether the CRC-CRN correctly predicts the effects of the mutation of a single gene, we have focused our attention on the mutations that are more common in CRC cancerogenesis (Fearon and Vogelstein, 1990; Smith et al., 2002), namely the GoF of k-Ras, and the LoF of APC, SMAD4, and p53. More specifically, for each one of the four mutations, we have separately computed its impact on the concentrations of the other proteins involved in the CRC network.

On the horizontal axis of Figure 2, we report the proteins *A*_*i*_, *i* = 1, …, 419, of the network. For each protein, on the vertical axis, we show the relative difference

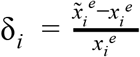

where 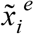 and 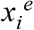 are the values of the concentration of *A*_*i*_ at the mutated and physiological equilibrium, respectively. A value of δ_*i*_ different from zero means that in the mutated network the concentration of the protein *A*_*i*_ is either increased (δ_*i*_ > 0) or reduced (δ_*i*_ < 0). In particular, a value of δ_*i*_ equal to −1 means that the function of protein *A*_*i*_ is almost completely stopped in the mutated network. In more general terms, the value of δ_*i*_ quantifies the relative change of the protein concentration, normalized by its value in the physiological network, and thus enables identifying which proteins are more sensitive to each one of the four considered mutations.

**Figure 2:**
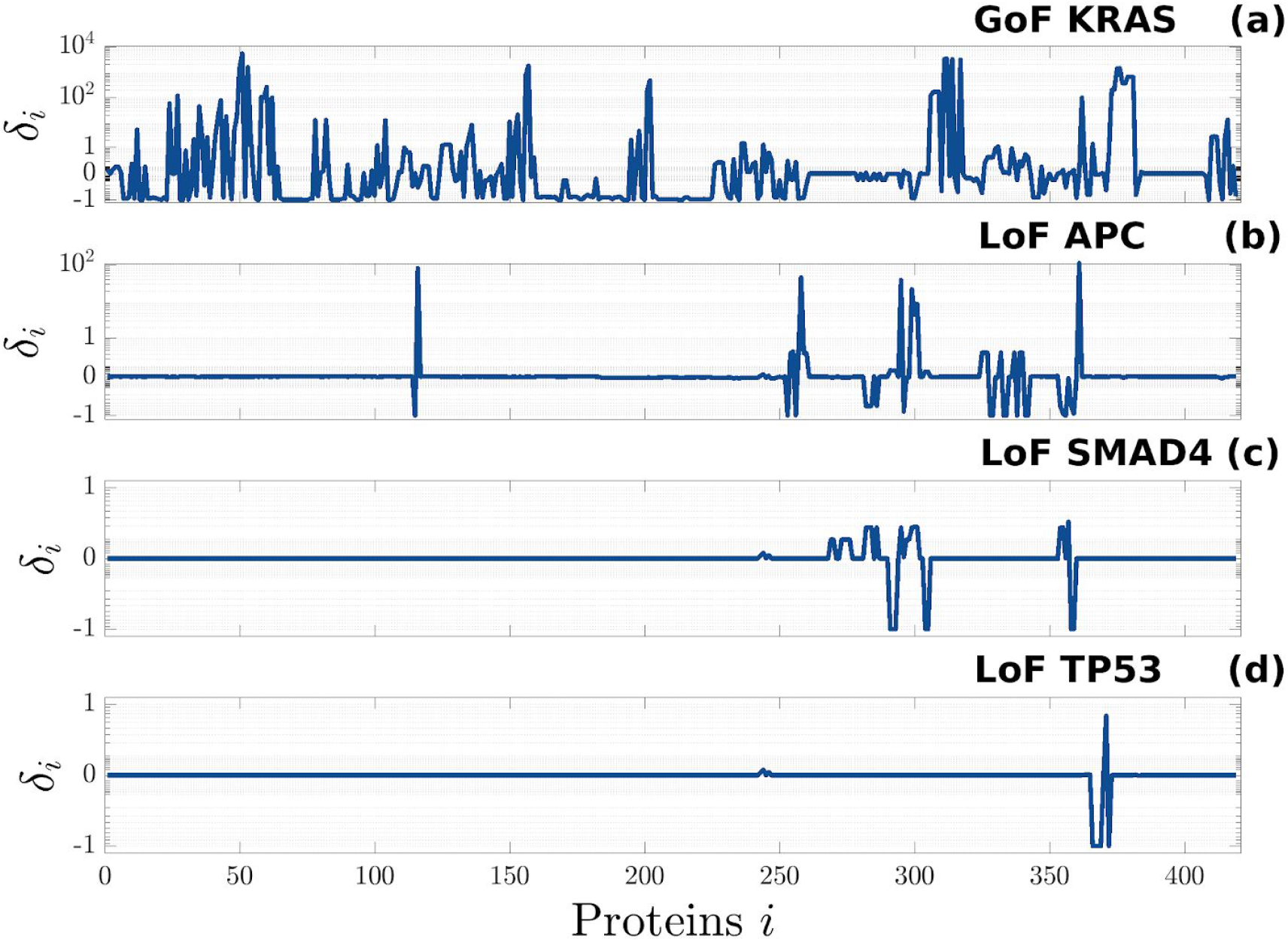
Relative difference between the concentrations at the equilibrium of the mutated and the physiological network. Each panel shows the result obtained by modifying the original CRC-CRN in order to simulate the effect of a different mutation, namely: (a) the GoF of KRAS, (b) the LoF of APC, (c) the LoF of SMAD4, (d) the LoF of TP53. For ease of visualization, on the horizontal axis we reported an index i = 1, …, 419. The name of the corresponding proteins *A*_*i*_ can be read in Table S1, while the list of the proteins significantly affected (i.e. | δ_*i*_ | > 0.03) by each one of the four mutations, together with the corresponding value of δ_*i*_ can be found in Table S2. For a complete explanation of the abbreviations used for the protein names we refer to (Tortolina et al. 2015)

The results shown in Figure 2, highlight how mutations may have a different impact on the CRC-CRN when acting on proteins at different levels of the network. Indeed, more than 76% of the proteins are significantly affected by a GoF of k-Ras, a protein upstream in the network, while about only 10.5%, 6.2%, and 1.4% of the proteins are affected by the LoF of APC, SMAD4, and TP53, respectively. Additionally, we found the strongest value of our index δ_*i*_ in correspondence with the GoF of k-Ras and the LoF of APC, Figure 2 (a) and (b) respectively. In the former case, a few proteins reach a value of δ_*i*_ close to 10^4^, as a consequence of the fact that they have a very low equilibrium concentration in the physiological network. Consider for example p-p-ERK: in the physiological cell 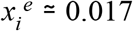, while in the network affected by a GoF of k-Ras, 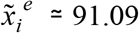 and thus δ_*i*_ ≃ 5, 37 · 10^3^.

These results suggest that, although KRAS, APC, SMAD4, and TP53 gene mutations are essential events for colorectal cancer development (Tsilimigras et al., 2018), the downstream effects of their mutation is more evident for the protein upstream in the signaling pathway. Moreover, literature reports that KRAS and APC mutations are the principal causes of the CRC onset, but they are not related to the tumor stage or location (Calistri et al., 2005; Fodde, 2002). Conversely, TP53 mutations seem to increase parallelly with the tumor stage, suggesting that this gene plays a pivotal role in the progression of CRC, more than in the pathology onset (Calistri et al., 2005). Regarding the role of SMAD4, it displays a pivotal role both in the development and in the progression of CRC (Mehrvarz Sarshekeh et al., 2017). Somatic mutations of SMAD4 are associated with more aggressive tumor biology, poor response to chemotherapy, metastases, and unfavorable overall survival among patients with resectable and unresectable CRC (Chung et al., 2018; Mizuno et al., 2018). On the other hand, the mutations of upstream proteins of a specific pathway could be counterbalanced by the activation/inhibition of other correlated pathways, while the mutation of downstream proteins, albeit of a minor entity, could determine major damages, since their activity could not be replaced by other pathways.

For each one of the four mutations, a comprehensive list of the proteins whose concentration significantly changes in the mutated network can be found in Table S2.

### GoF of PI3K, k-Ras and Raf, and LoF of PTEN and AKT determine an alteration of the p53 level

As in the previous subsection, we considered a set of single-gene mutations. The new specific aim is to compare the effects of each mutation on the same target molecule, in order to show how the CRC-CRN can be used to highlight mechanisms that can alter the function of a given protein. In view of its connection with CRC, we focused on p53.

In the previous section, we considered a mutation resulting in the LoF of TP53 and we quantified the alteration induced by such a mutation on the values of the concentrations of all the proteins within our CRC-CRN. As shown in Figure 2 (d), since p53 is a downstream protein in our network, the LoF of TP53 alters the concentration of only a few proteins. On the contrary, even when no mutation directly involves the gene TP53, the value of the equilibrium concentration of the protein p53 may be altered by various mutations affecting other proteins located at an upstream level of the CRN (Rivlin et al., 2011). Motivated by these considerations, here we assume that TP53 is not affected by any mutation, and we show how to use the proposed tool to infer the mechanisms altering the concentration of p53 as a consequence of an upstream mutation.

To this end, after selecting a set of mutations to be tested, we computed the equilibrium of the corresponding mutated network and we calculated the relative difference δ_*p*53_ between the concentration of p53 in the mutated and the physiological equilibrium. Table 1 shows the value of δ_*p*53_ for a set of mutations that significantly impact the concentration of the protein. In detail, the concentration of p53 was reduced by about 0.7 times the value in the physiological network by both the GoF of PI3K and the LoF of PTEN, while it was increased by the GoF of k-Ras and the GoF of Raf. However, the strongest effect is induced by the LoF of AKT in which case the value of δ_*p*53_ was found equal to 130.5.

**Table 1:** Relative difference δ_*p*53_ of the equilibrium concentrations of p53 induced by a set of single-gene mutations. The first and second rows report the considered mutation and the corresponding value of δ_*p*53_, respectively.

| Mutation | GoF PI3K | LoF PTEN | GoF KRAS | GoF BRAF | LoF AKT |
| --- | --- | --- | --- | --- | --- |
| $\delta_{p53}$ | -0.66 | -0.68 | 1.38 | 2.28 | 130.5 |

To understand the reasons underlying these alterations, we observe that in our CRC-CRN the degradation of p53 is regulated by the phosphorylated form of MDM2, whose activation is in turn regulated by phospho-AKT (p-AKT) (Li and Kurokawa, 2015). Therefore, in Figure 3, we show the concentration of MDM2, AKT, and their phosphorylated form p-MDM2 and p-AKT in the physiological network (panel (a)) and the altered values induced by the 5 mutations mentioned in Table 1 (panel (b)-(f)). We observe that both the GoF of PI3K and the LoF of PTEN promote the phosphorylation of AKT and MDM2 thus speeding the degradation of p53. On the other hand, it was demonstrated on CRC cellular models that activation of the PI3K/AKT pathway inhibits the apoptosis, cell growth, and modulation of cellular metabolism, lowering p53 and PTEN concentration (De Roock et al., 2011; Liu et al., 2020). Conversely, the activation of the PTEN pathway decreases the PI3K/AKT-dependent cellular proliferation and regulates the stability of p53 (De Roock et al., 2011; Georgescu, 2010).

**Figure 3:**
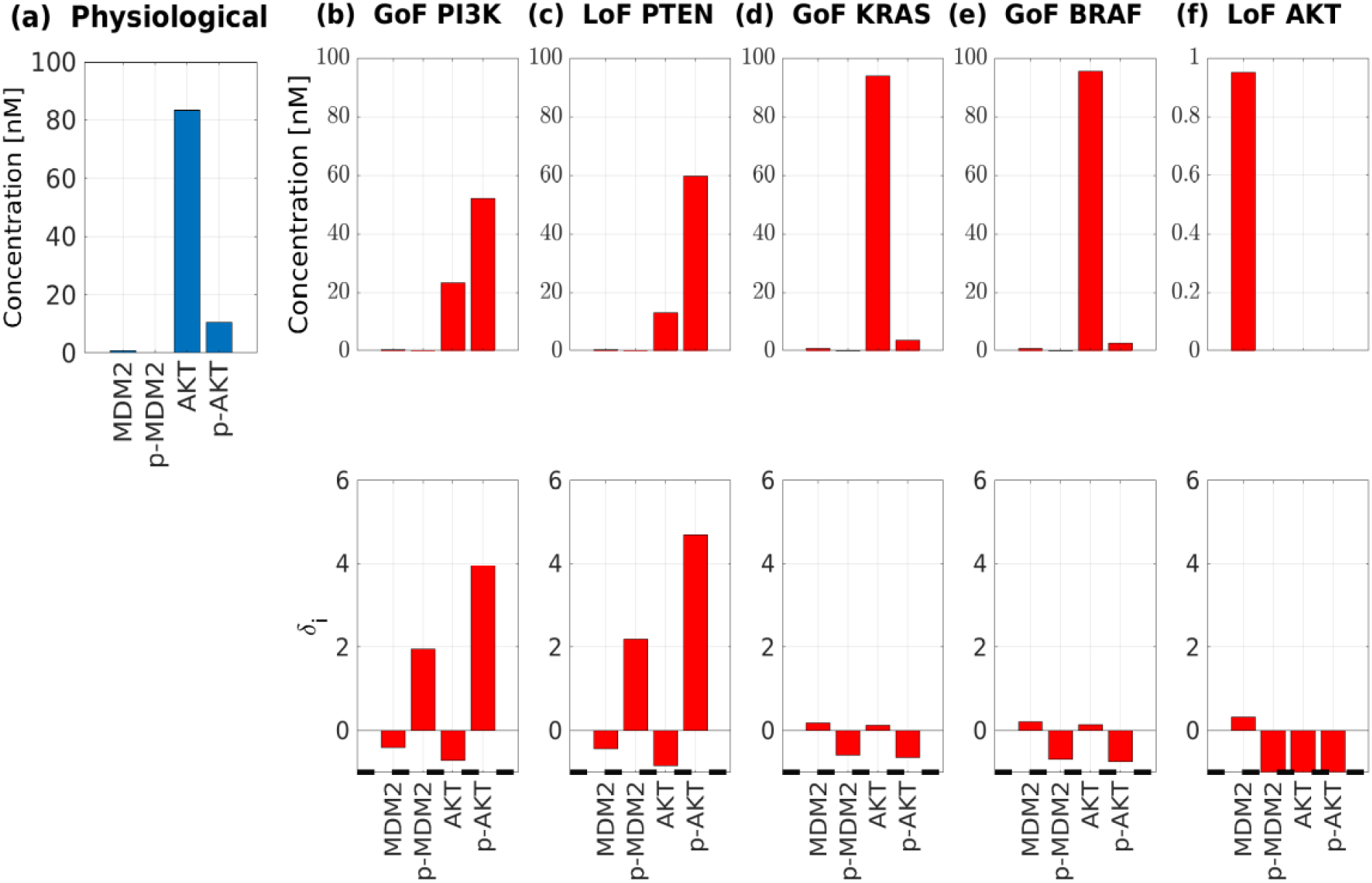
Effects of various single-gene mutations on the value of p53 concentration. (a) Value of the concentrations at the equilibrium of the physiological network of the proteins (directly) involved in the degradation of p53, namely MDM2, AKT, and their phosphorylated form p-MDM2 and p-AKT, respectively. (b)-(f) Effect on these concentrations of 5 mutations, namely GoF of PI3K, LoF of PTEN, GoF of KRAS, GoF of BRAF, and LoF of AKT. For each mutation in the upper panel, we show the values of the concentrations at the equilibrium of the mutated network. In the lower panel, the corresponding relative difference δ_*i*_ is depicted.

Moreover, our model shows that the GoF of KRAS, and the GoF of BRAF (Figure 3 (d)-(e)), downregulate the phosphorylation of AKT, which, in turn, determines the increment of the p53 level. When the LoF of AKT is considered, the phosphorylation of AKT and MDM2 is completely stopped. Indeed, as shown in Figure 3 (f) in this case the value of δ_*i*_ for p-MDM2 and p-AKT is equal to −1. This explains why in all these mutations the concentration of p53 is increased, but the strongest effect is induced by the LoF of AKT, confirming the relation between RAS/RAF/MEK/ERK signaling axis, the inhibition of AKT pathway, and the intracellular concentration of p53 (Fritsche-Guenther et al., 2016).

None of the considered mutated proteins is related to p53 by a specific, direct chemical reaction. Furthermore, the proteins are rather far from p53 in the network topology and do not belong to the same pathway. Nevertheless, the proposed network approach has disclosed their indirect influence on p53.

### Multiple-gene mutations: effects of two mutations combination on p53 level

Most cancers develop following the accumulation of a series of specific mutations in the cell. Thus, characterizing the impact of the interaction among a group of mutations plays a crucial role in the prediction of tumor progression. Quantifying the combined effect of a set of mutations is not trivial, also when the effects of each single mutation are known, because some mutations may actually induce opposite effects on a given protein. For example, Table 1 shows that the GoF of PI3K and the LoF of PTEN reduce the concentration of p53, while the GoF of k-Ras, the GoF of Raf, and the LoF of AKT individually increase it.

The proposed tool easily allows one to overcome this problem. Indeed, as described in the Model and Methods section, the simultaneous action of a group of mutations can be simulated by changing the initial conditions of the system according to each LoF of the group and by removing the reactions associated with each GoF. To show an application, Table 2 synthesizes the results obtained when 6 pairs of mutations are considered, each composed of a mutation that downregulates p53 and a mutation that instead augments its concentration. We observe that when the GoF of PI3K is combined with either the GoF of k-Ras or the GoF of Raf, the concentration of p53 decreases. This may depend on the negative control exerted by the Ras/Raf/MEK/ERK pathway on the downstream AKT signal (Fritsche-Guenther et al., 2016).

**Table 2.** The relative difference δ_*p*53_ of the equilibrium concentrations of p53 induced by pairs of simultaneous mutations. Each element of the table shows the value of δ_*p*53_ obtained when the mutations reported in the corresponding row and column are incorporated in the network.

|  | GoF KRAS | GoF BRAF | LoF AKT |
| --- | --- | --- | --- |
| GoF PI3K | -0.65 | -0.65 | 130.49 |
| LoF PTEN | -0.68 | -0.68 | 130.49 |

On the contrary, the LoF of AKT prevails on the GoF of PI3K as their combination increases p53 concentration. This could be due to the relationship between PI3K and AKT since PI3K activation is upstream to the AKT signal (Naderali et al., 2019). Therefore, AKT reduction plays a predominant role on p53 levels regardless of PI3K activation. Analogous results hold for the LoF of PTEN.

Interestingly, the values obtained for the GoF of KRAS and BRAF are the same probably because the proteins codified by these two genes belong to the same pathway, and, in detail, the second is downstream of the first. The same observation can be made also for the GoF of PI3K and the LoF of PTEN.

### Effect of Dabrafenib on the CRC-CRN

Assuming that the CRC-CRN is affected by a mutation resulting in the GoF of k-Ras, we investigated the effect on the mutated network of Dabrafenib, a drug that inhibits Ras activity (Morkel et al., 2015; Puszkiel et al., 2019).

To this end, we modelled the drug as a competitive inhibitor (Ingalls, 2013), and we added to the CRC-CRN the reversible reaction

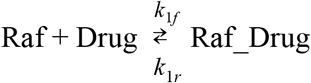

where DRUG stands for the considered drug, in our example Dabrafenib, and Raf_DRUG is the inactive drug-target complex. Additionally, we assume that *k*_1*f*_ ≫ *k*_1*r*_ so that the drug binds almost steadily to the targeted molecules. Specifically, in our simulation we set *k*_1*f*_ = 0.5 (nM s) ^−1^, which is the average rate constant over all the second-order reactions within the CRC-CRN, and *k*_1*r*_ = 0.005*s*^−1^. Provided that the ratio between the two rate constants is kept fixed, different values of *k*_1*f*_ and *k*_1*r*_ will impact the speed at which the modified CRC-CRN reaches the equilibrium but will not significantly alter the final concentration profile.

We then quantified the effect of the drug delivery on the protein concentrations as follows. We modified the CRC-CRN by accounting for both the GoF of k-Ras and the action of the drug targeting Raf. We then integrated the corresponding system of ODEs with initial values of the proteins concentrations equal to the values of the steady-state of the network affected by GoF of k-Ras, and with different values of the drug initial concentration. Specifically, we set

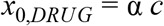

Where *c* = 50 nM is the total molar concentration at disposal of the proteins within Raf conservation law and α ∈ {1, 0.75, 0.5, 0.25}.

Figure 4 shows the changes induced by the drug on the whole equilibrium concentration profile. We observe that the drug reaches its highest effectiveness when α = 0.75 i.e. when about 37 nM of the drug is delivered to the cell. In this case, the concentrations of most of the proteins involved in the cellular signaling of CRC return to a value close to the steady-state of the physiological condition. Only for a few proteins, the value of the concentration remains different from the physiological one, even after the drug action. These are: (i) Raf, whose function is correctly inhibited; (ii) a group of complexes that involve the activated form of k-Ras, that is still overexpressed; (iii) the complexes that are products of the reactions removed to simulate the GoF of k-Ras, whose function is thus stopped, and a group of corresponding complexes, whose concentration increases so that the total molar concentration within the conservation laws is conserved.

**Figure 4:**
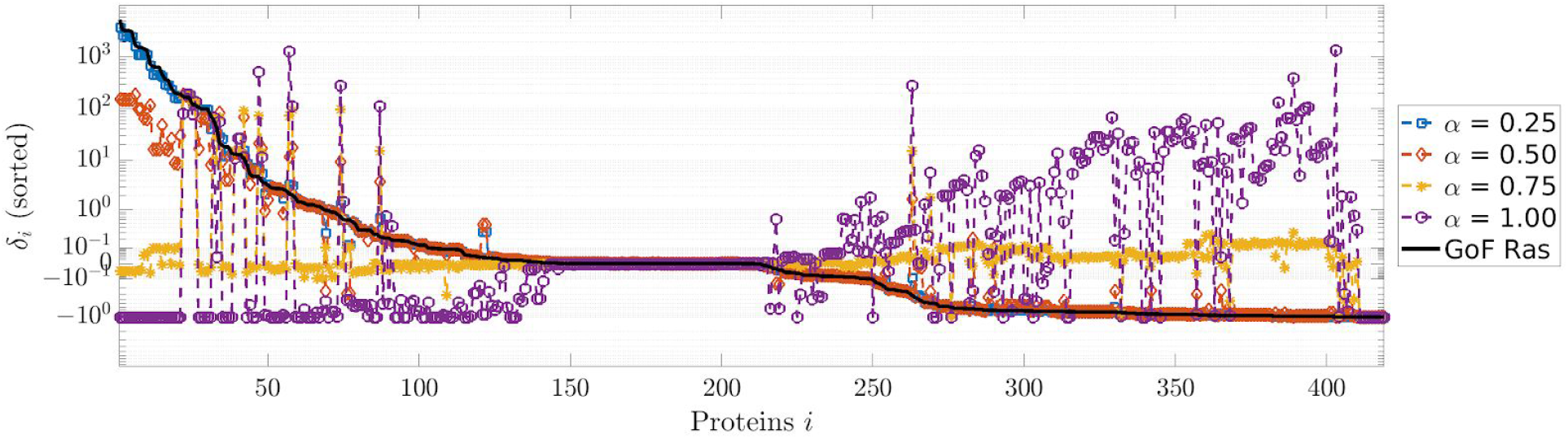
Effects of the drug on the whole protein concentrations profile. The black line represents the relative difference between the concentrations at the equilibrium of the network affected by a GoF of k-Ras and the concentrations at the physiological equilibrium; all the values are sorted in decreasing order. The values δ_*i*_ for all the proteins can be found in Table S5, second column. The colored lines represent the relative difference between the equilibrium concentrations obtained after that the drug has been incorporated in the mutated network and the concentrations in the physiological equilibrium. Each color corresponds to a different value of the initial drug concentration, parameterized by found in Table S5, third column. α. The values of δ_*i*_ when α = 0.75 can be found in Table S5, third column.

Our approach also enabled us to quantify the effect of an under- or over-dosed administration. Indeed, Figure 4 shows that when *x*_0,*DRUG*_ is too small, the drug essentially has no impact on the values of the proteins concentrations, while a too high value may result in severe side effects.

To better understand the mechanism underlying the described network response, we focused on the Ras-Raf-MEK-ERK cascade, i.e. a pathway involving Raf, which is the target of the simulated drug. Acting as a competitive inhibitor, the drug binds Raf to the inactive complex Raf_DRUG. As shown in Figure 5 (b), this results in a fast decrease of the concentrations of both Raf and its phosphorylated form p-Raf. Interestingly, when α = 0.75, p-Raf reaches an equilibrium value equal to that of the physiological network. Shortly thereafter, the reduction of p-Raf concentration downregulates the phosphorylation of MEK and ERK. Indeed, Figure 5 (b) shows an increase in the concentration of both the proteins at the expense of their phosphorylated form p-MEK and p-ERK. The inhibition of Raf also affects the proteins upstream in the network. For example, as shown in Figure 5 (b), the concentration of Ras increases as a result of the drug action, probably as an attempt to induce the activation of Raf. On the other hand, these data based on our model are confirmed by the literature data, which show that the Dabrafenib treatment on CRC cellular model, by inhibiting the Raf activation, increases the Ras level and diminishes the MAPK pathway (Haarberg and Smalley, 2014).

**Figure 5:**
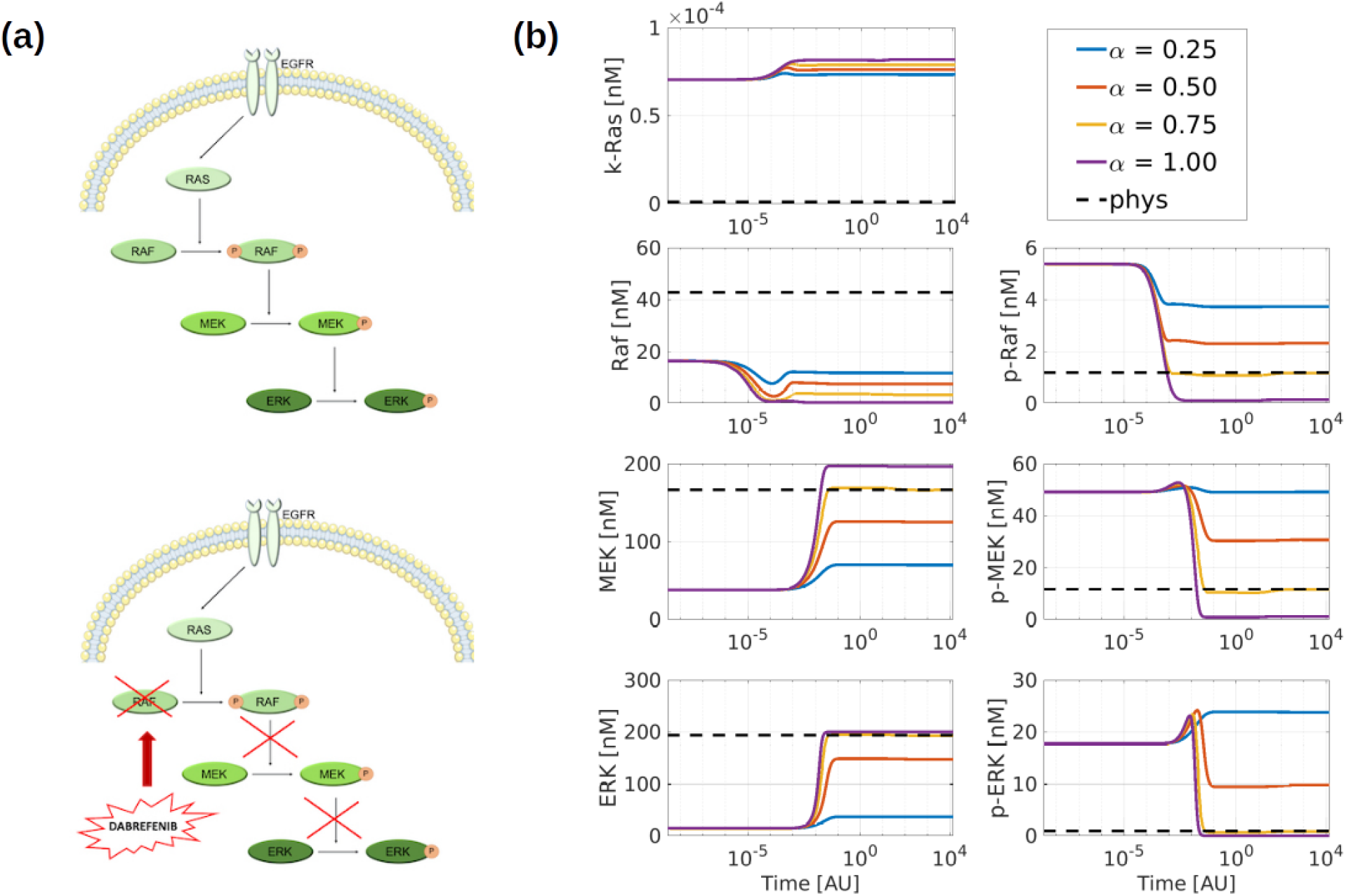
Effects of the drug on the Ras-Raf-MEK-ERK cascade (MAPK pathway). (a) Schematic representation of the MAPK pathway extracted from the CRC-CRN and of the changes induced by the drug Dabrafenib modeled as a competitive inhibitor of Raf kinase. (b) Time-courses of the concentrations of the proteins within the MAPK cascade obtained by solving the system of ODEs associated to the modified CRC-CRN with the initial state set equal to the equilibrium of the network affected by a GoF mutation of KRAS. Different lines correspond to a different value of the initial concentration of the drug; the black dotted line depicts the value of the protein concentrations in the physiological cell that is taken as reference.

To unravel the mechanism underlying this feedback effect, in Figure 6 we plot the fluxes of the reactions involving Ras and its active form Ras_GTP as functions of time. For the reversible reactions, the sum of the forward and backward fluxes is considered. In Figure 6 (c)-(d) we report the fluxes that are significantly different from zero and the corresponding chemical reactions. The Figure shows that the network reacts to the reduction of p-Raf concentration by decomposing the complex Raf_Ras_GTP into p-Raf and Ras_GTP. This causes an increase of Ras_GTP concentration that in turns promotes the production of Ras through the decomposition Ras_GTP → Ras + GTP.

**Figure 6.**
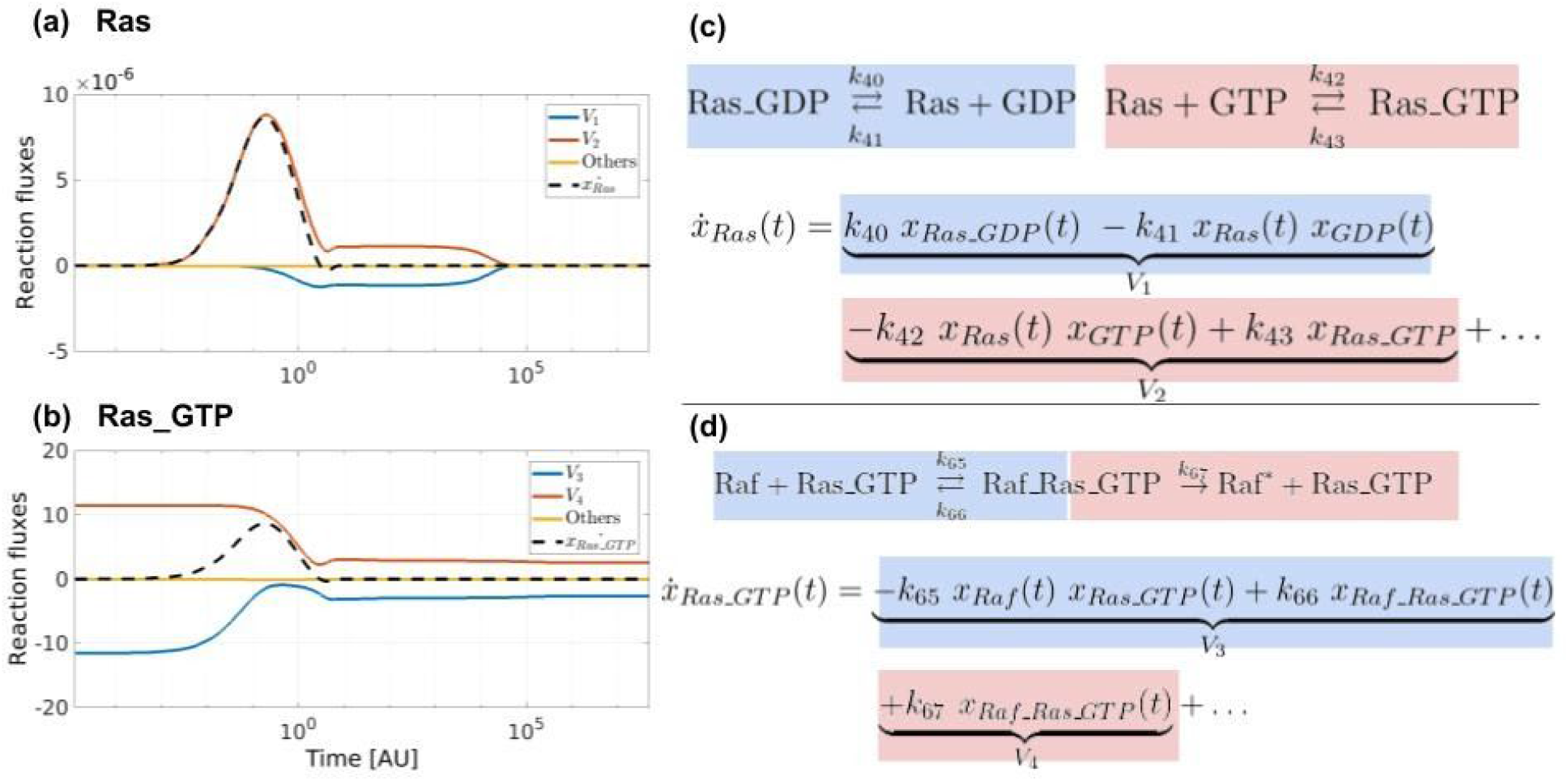
Analysis of the flux rates of the reactions involving Ras and Ras_GTP. (a-b) Flux rates that contribute to the dynamics of the concentrations of Ras (inactivated form) and Ras_GTP (the activated form) respectively. (c-d) Mathematical expression of the derivatives of such proteins and corresponding chemical reactions.

As a final remark, we point out that the concentrations of Ras and Raf start changing right after the delivery of the drug and reach an equilibrium value in a rather small time interval. Changes in MEK and ERK concentrations are simultaneous between each other and occur sometime after that k-Ras and Raf reach equilibrium. Since Ras is located upward of Raf in the pathway shown in Fig 5 (a), the change in Raf may be regarded as a feedback effect. Changes in MEK and ERK, which occur after Raf has reached equilibrium, may be associated with delay effects.

## Discussion

In this work, we have shown how a computational tool for simulating signal transduction networks can be applied for modeling the information flow inside a CRC cell at the G1-S transition point. Results reported in this work show that the proposed model based on a whole molecular interaction map displays several advantages in evaluating the changes in CRC cell signaling induced by mutations and drugs, over classical single-pathway approaches.

First of all, with the proposed mathematical model we are able to quantify the global effects induced on the whole network by local changes due to the mutation of one or more genes. In detail, we considered two particular classes of mutations that result in either the loss or the gain of function of specific proteins of the network. By exploiting this feature, we described the alterations induced on the concentrations of all the proteins within the network by the four mutations, more commonly found in CRC cells, namely the LoF of APC, SMAD4, and p53 end the GoF of k-Ras. For each one of the considered mutations, we computed and analyzed the equilibrium states of the physiological and mutated network. In particular, we introduced the relative difference between mutated and physiological equilibrium values of the protein concentrations as a quantitative index to identify which proteins are affected by each mutation, but also to quantify the strength of such effects.

By simultaneously analyzing several interconnected pathways, our tool also allowed us to highlight links between the different proteins that would not be evident by studying a single pathway at a time. Additionally, our framework can be easily adapted to simulate the effect of the occurrence of multiple mutations. As an example, we considered TP53, a gene that has proved to play a pivotal role in CRC progression. In fact, even when TP53 is not directly affected by any mutations, alterations of proteins upstream in the network may induce changes in the concentration of the protein p53. In this work, we quantified the changes induced on the concentration of unmutated p53 by 5 mutations, namely GoF of PI3K, GoF of KRAS, GoF of BRAF, LoF of PTEN, and LoF of AKT, considered one at the time or coupled to simulate the action of two concurrent mutations.

Eventually, we employed our tool to simulate the action of targeted drugs on the CRC cells signaling. In detail, we considered a network affected by a GoF mutation of KRAS, and we analyzed the action of Dabrafenib, modeled as a competitive inhibitor targeting BRAF kinase. By looking at the changes induced on the whole protein concentrations profile, we were able to: (i) obtain a detailed description of the action of the drug on the MAPK pathway, as well as on the other elements of the network; (ii) identify an amount of the drug (37 nM in our simulation) capable of restoring a value of most of the protein concentrations close to that in the physiological network; (iii) propose a reasonable interpretation of the results, in terms of time courses of reaction fluxes.

Although preliminary, these results show that the proposed method is capable of predicting the quantitative effects of targeted drugs and thus could represent a valuable support in the design and optimization of novel targeted therapies. In this work, we limited our attention to a kinase inhibitor acting on CRC cells. Future efforts will be devoted to extending the proposed model to different types of drug and cancer cells, and to investigate the interplay between cytoplasmic protein alterations and genomic mutations in order to supply a more comprehensive model of different types of LoF and GoF mutations, including a wider class of mechanisms altering the protein function, such as e.g. copy number variations. Finally, the results of the presented paper have been validated by using literature papers. A more systematic validation through properly designed biological experiments is our next goal.

## Model and Methods

### A mathematical model for CRNs

Our CRC-CRN models the G1-S transition in HCT116 and HT29 CRC cell lines as a complex CRN describing the flow of information through 10 interacting pathways (Tortolina et al., 2015). A total of r=850 reactions involving n=419 well-mix proteins were included in our network. The list of all the considered proteins and chemical reactions can be found in Table S1 and Table S3, respectively.

By applying the law of mass action, the kinetic of the proteins concentrations can be modeled through a system of n ordinary differential equations (ODEs) of the form (Chellaboina et al., 2009; Feinberg, 1987; Yu and Craciun, 2018)

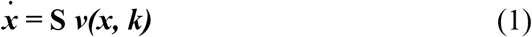

where *x* = (*x*_1_, …, *x_n_*)^*T*^ is the state vector identified by the molar concentrations (nM) of the proteins, the superimposed dot denotes the time-derivative, **S** is the constant stoichiometric matrix of size n x r, ***v(x, k)*** denotes the time-variant vector of reaction fluxes of length r, and ***k*** = (*k*_1_, …, *k_r_*)^*T*^ is the set of known reaction rate constants, whose value can be found in Table S3. In equation (1) we have assumed that all the molecular exchanges between the cell and the environment are encoded in the stoichiometric matrix.

In this work, we were mainly interested in characterizing the state of the system when the network reaches equilibrium. To this end, after setting the initial values of the protein concentrations, we integrate the system of ODEs (1), using the Matlab tool ode15s (Shampine and Reichelt, 1997), and we took the asymptotic value of the obtained solution as the equilibrium point.

Our model of LoF and GoF mutations built on the analysis of the moiety conservation laws of the system. Each conservation law identifies a group of proteins whose aggregate concentrations do not change with time and is formally defined as a set of positive, integer coefficients **γ**= (γ_1_, …, γ_n_)^T^ such that the product **γ**^T^ ***x****(t)* remains fixed over the simulated concentrations dynamic. A set of generators for all the moiety conservation laws of the system can be computed by studying the left null space of the stoichiometric matrix (De Martino et al., 2014; Schuster and Höfer, 1991). By applying this procedure to the CRC-CRN we obtained 81 independent moiety conservation laws involving all the proteins within the network but 10. The latter are either proteins that undergo degradation or proteins that have direct contact with the external environment.

The importance of conservation laws is twofold. On the one hand, we have numerically verified (Sommariva et al., 2020) that the system of ODEs (1) admits a unique equilibrium on the set

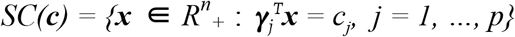

where **γ**_1_, …, **γ**_p_ are the p=81 independent constant generators of the moiety conservation laws and ***c*** = (*c*_1_, …, *c_p_*) is the vector of the corresponding constant aggregation concentrations. The set *SC(**c**)* is called the stoichiometric compatibility class. On the other hand, we made use of the conservation laws to simplify the input parameters required by our simulation tool for computing the equilibrium states. In fact, the state of the network, mimicking either a physiological or a mutated cell, is fully characterized by assigning the set **c**of the constant aggregate concentrations. Indeed, the value of ***c*** defines a unique stoichiometric compatibility class and all initial states belonging to the same class lead to the same stationary state. Therefore, every equilibrium state corresponds to a set of 81 constants, and conversely (Sommariva et al., 2020).

### LoF mutations

By exploiting the mathematical framework developed in our previous work (Sommariva et al., 2020), we simulated the effect of a LoF of APC, AKT, SMAD4, and PTEN by observing that all these proteins are involved in only one conservation law. Their LoF mutations are implemented by setting to zero the total concentration at disposal of the corresponding conservation law. In practice, this is achieved by projecting the initial concentration values describing the physiological cell into a new initial state where the concentrations of the mutated protein and of all its compounds are set to zero.

In this work, we also simulated the LoF of p53 by downregulating its production. Since p53 undergoes degradation, it is not involved in any concentration law and thus the previous framework does not hold. However, in our MIM the production of p53 is modeled by the presence of an auxiliary variable, called *p53_generator*, whose concentration is assumed to be constant to model the presence of a pool that constantly feeds the production of p53. In the mutated cell, such production is stopped by setting to zero the concentration of *p53_generator*. Both the described procedures simulate the effect of null mutations where the function of the mutated proteins is totally lost and the concentrations of the related molecules vanish. However, mutation with a different degree of loss of function can be easily achieved by setting the amount of available total concentration to a value lower than the one in the physiological cell but different from zero.

### GoF mutations

In this work, we quantified the effect of mutations resulting in the GoF of k-Ras, Raf, PI3K, and Betacatenin. The GoF of a given protein is induced by removing from the CRC-CRN all the reactions involved in its deactivation.

As an example, the dephosphorylation of Raf is modeled in our CRC-CRN through the set of reactions

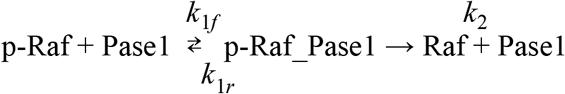

where p-Raf is the activated form of Raf, consisting in the phosphorylation of a specific amino acid.

A GoF of Raf is induced by setting to zero the rate constants in the two forward reactions, namely, we set *k*_1*f*_ = *k*_2_ = 0. This results in a novel set of n ODEs

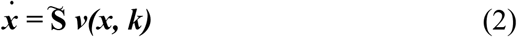

where the stoichiometric matrix 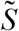 is defined by setting to zero the two columns of the matrix **S** in (1) corresponding to the deleted chemical reactions. By removing only the two forward reactions, the deactivation of p-Raf is completely blocked while the rank of the novel stoichiometric matrix is kept equal to that of the original stoichiometric matrix. As a consequence, the reduced system of ODEs described by equation (2) maintains the same conservation laws of the original system (1).

The list of reactions removed when implementing each one of the GoF mutations considered in this paper, namely the GoF of BRAF, the GoF of k-Ras, and the GoF of PI3K, can be found in Table S4.

Similarly to what we did for the LoF mutations, mutations resulting in different degrees of gain of function of the considered protein, can be modeled by reducing the value of the rate constants *k*_1*f*_ and *k*_2_. By doing so, the deactivation of Raf still takes place, but at a slower speed than in the physiological cell.

### Combination of multiple LoF and GoF mutations

Consider a cell affected by a number of mutations each one of them resulting in the loss or gain of function of a specific protein. As we have shown in a previous work (Sommariva et al., 2020), our framework allows us quantifying the simultaneous effect of all mutations.

Specifically, each LoF mutation entails a change in the value of the total concentrations provided to the algorithm. Instead, to account for the GoF mutations we modify system (1) by reducing or zeroing the value of the rate constants corresponding to reactions involved in the deactivation of the proteins affected by this type of mutations. The value of the protein concentrations at the equilibrium is computed by integrating the modified system of ODEs with initial conditions defined according to the constraints imposed by the LoF mutations.

Importantly, the same set of steady-state values would have been obtained by starting from the system modeling the physiological state of the cell and iteratively computing the equilibrium of the modified system accounting for an increasing number of mutations, regardless of their order.

## Acknowledgment

S.S. kindly acknowledges the AIRC grant COENzYME: “Chemotherapy effect On cell ENergY Metabolism and Endoplasmic reticulum redox control”.

## Author contribution

Project conceptualization: G.C., F.B., L.T, N.C., S.P, and M.P.. Software: S.S. Formal analysis and visualization: S.S., G.C, S.R., and M.P.. Writing - original draft: S.S., G.C., S.R., F.F. and M.P. Writing - review & editing: all authors.

## Supplemental Information

Table S1. Table of the proteins *A*_*i*_, *i* = 1, …, 419, involved in the CRC-CRN related to Figure 2. Also the values of the initial concentrations used for computing the equilibrium state for the healthy cell are reported.

File: table_S1.xlsx

Table S2. Table of the proteins significantly affected (i.e. | δ_*i*_ | > 0.03) by the mutations related to Figure 2.

File: table_S2.xlsx

Table S3. Table of the chemical reactions and of the values of the rate constants involved in the CRC-CRN related to the Model and Methods section, paragraph A mathematical model for CRNs.

File: table_S3.xlsx

Table S4. Table of the chemical reactions removed when implemented each of the GoF mutations related to Model and Methods section, paragraph GoF mutations.

File: table_S4.xlsx

Table S5. Table listing (i) the relative difference between the concentrations at the equilibrium of the network affected by a GoF of k-Ras and the concentrations at the physiological equilibrium and (ii) the relative difference between the equilibrium concentrations obtained after that about 37 nM of drug has been incorporated in the mutated network and the concentrations in the physiological equilibrium, related to Figure 4.

File: table_S5.xlsx

